## Supplementary Material, Tables and figures for "Optimal microRNA sequencing depth to predict cancer patient survival with random forest and Cox models"

### Supplementary Materials

#### 1 More details on the Cox model

The function $L$ is called the 'pseudo-likelihood', because it is not a product of density functions, but a product of conditional probabilities. $\hat{\boldsymbol{\beta}}$ is computed by maximizing this pseudo-likelihood function: $\hat{\boldsymbol{\beta}} = \arg\max_{\boldsymbol{\beta}}(l(\boldsymbol{\beta}))$, with $l(\boldsymbol{\beta}) = \log(L(\boldsymbol{\beta}))$, the log-pseudo-likelihood.

Note that the Cox model is not intuitive, in the sense that it links genetic data to patient survival in an indirect way, through the hazard function. However, Cox pseudo-likelihood allows censored data to be efficiently dealt with. Moreover, this yields a robust inference procedure where the baseline function $h_0(t)$ does not need to be modeled or estimated in a parametric way. Finally, the estimation procedure leads to a convex optimization problem, for which efficient procedures and packages exist for computing $\hat{\boldsymbol{\beta}}$ [Friedman et al., 2010].

#### 2 More details on the penalization methods

The $\ell_1$ norm forces some coefficient estimates $\hat{\beta}_j, j = 1, ..., p$ to be zero, and allows the selection to be made. For multivariate Cox selection models, the genes selected are defined as the genes with nonzero $\hat{\beta}_j$ coefficients. It has been empirically observed that if there are high correlations between predictors, the ridge penalty provides better prediction performance than the lasso [Tibshirani, 1997]. The elastic net penalty have been developed to tackle this issue.

We computed the weight of the penalty, $\lambda$, by K-fold cross-validation (K = 5) using the R package *glmnet* [Friedman et al., 2010]. The weight $\lambda$ that minimizes deviation in the cross-validation is given by $\lambda_{min}$. We chose $\alpha = 0.3$ in the elastic net, as the deviance remains stable for different values of $\alpha$ and the number of genes selected starts to stabilize below this value (Supplementary Fig. S1).

For more details of the mathematical concepts used in this article, we refer the reader to the book 'The Statistical Analysis of Failure Time Data' [Kalbfleisch and Prentice, 2011].

### Supplementary Tables

| Cancer | Name |
|---|---|
| LAML | Acute Myeloid Leukemia |
| ACC | Adrenocortical carcinoma |
| BLCA | Bladder Urothelial Carcinoma |
| LGG | Brain Lower Grade Glioma |
| BRCA | Breast invasive carcinoma |
| CESC | Cervical squamous cell carcinoma and endocervical adenocarcinoma |
| CHOL | Cholangiocarcinoma |
| LCML | Chronic Myelogenous Leukemia |
| COAD | Colon adenocarcinoma |
| ESCA | Esophageal carcinoma |
| GBM | Glioblastoma multiforme |
| HNSC | Head and Neck squamous cell carcinoma |
| KICH | Kidney Chromophobe |
| KIRC | Kidney renal clear cell carcinoma |
| KIRP | Kidney renal papillary cell carcinoma |
| LIHC | Liver hepatocellular carcinoma |
| LUAD | Lung adenocarcinoma |
| LUSC | Lung squamous cell carcinoma |
| DLBC | Lymphoid Neoplasm Diffuse Large B-cell Lymphoma |
| MESO | Mesothelioma |
| MISC | Miscellaneous |
| OV | Ovarian serous cystadenocarcinoma |
| PAAD | Pancreatic adenocarcinoma |
| PCPG | Pheochromocytoma and Paraganglioma |
| PRAD | Prostate adenocarcinoma |
| READ | Rectum adenocarcinoma |
| SARC | Sarcoma |
| SKCM | Skin Cutaneous Melanoma |
| STAD | Stomach adenocarcinoma |
| TGCT | Testicular Germ Cell Tumors |
| THYM | Thymoma |
| THCA | Thyroid carcinoma |
| UCS | Uterine Carcinosarcoma |
| UCEC | Uterine Corpus Endometrial Carcinoma |
| UVM | Uveal Melanoma |

**Supplementary Tab. S1. Acronym of the TCGA cancers**

**A**

| Cancer | UVM | ACC | KIRP | MESO | KIRC | LGG | CESC | LIHC | PRAD | LUAD | UCEC |
|---|---|---|---|---|---|---|---|---|---|---|---|
| Fold reduction | 100 | 10 | 5 | 5 | 10 | 10 | 1000 | 100 | 10 | 10 | 100 |
| Corresponding median sequencing depth | 50 | 500 | 1000 | 1000 | 200 | 700 | 5 | 50 | 400 | 500 | 40 |
| Metric degraded first | C-index | C-index | C-index | both | both | IBS | C-index | both | C-index | C-index | C-index |

**B**

| Cancer | UVM | ACC | KIRP | MESO | KIRC | LGG | CESC | LIHC | PRAD | LUAD | UCEC |
|---|---|---|---|---|---|---|---|---|---|---|---|
| Fold reduction | 100 | 100 | 100 | 100 | 1000 | 1000 | 100 | 100 | 100 | 100 | 100 |
| Corresponding median sequencing depth | 400 | 400 | 400 | 500 | 50 | 50 | 500 | 500 | 500 | 400 | 200 |
| Metric degraded first | IBS | both | C-index | both | both | both | C-index | C-index | C-index | C-index | C-index |

**Supplementary Tab. S2. Maximum fold reduction without degradation of the C-index and the IBS, corresponding median sequencing depth (thousands of reads), and prediction metric degraded first for random survival forest and for miRNA-seq data (A) and mRNA-seq data (B) for the 11 investigated cancers.**

**A** miRNA - Cox

| Cancer | UVM | ACC | KIRP | MESO | KIRC | LGG | CESC | LIHC | PRAD | LUAD | UCEC |
|---|---|---|---|---|---|---|---|---|---|---|---|
| Proportion of patients in the training set | 0.4 | 0.6 | 0.6 | 0.6 | 0.7 | 0.7 | 0.6 | 0.7 | 0.8 | 0.8 | 0.4 |
| Corresponding number of patients | 31 | 46 | 161 | 51 | 356 | 354 | 173 | 248 | 389 | 386 | 213 |

**B** miRNA – random survival forest

| Cancer | UVM | ACC | KIRP | MESO | KIRC | LGG | CESC | LIHC | PRAD | LUAD | UCEC |
|---|---|---|---|---|---|---|---|---|---|---|---|
| Proportion of patients in the training set | 0.3 | 0.4 | 0.2 | 0.5 | 0.4 | 0.5 | 0.1 | 0.5 | 0.3 | 0.5 | 0.2 |
| Corresponding number of patients | 23 | 31 | 54 | 42 | 203 | 253 | 29 | 178 | 146 | 242 | 106 |

**C** mRNA - Cox

| Cancer | UVM | ACC | KIRP | MESO | KIRC | LGG | CESC | LIHC | PRAD | LUAD | UCEC |
|---|---|---|---|---|---|---|---|---|---|---|---|
| Proportion of patients in the training set | 0.8 | 0.5 | 0.6 | 0.7 | 0.6 | 0.6 | 0.8 | 0.7 | 0.8 | 0.8 | 0.7 |
| Corresponding number of patients | 62 | 38 | 161 | 59 | 305 | 304 | 230 | 248 | 389 | 386 | 372 |

**D** mRNA – random survival forest

| Cancer | UVM | ACC | KIRP | MESO | KIRC | LGG | CESC | LIHC | PRAD | LUAD | UCEC |
|---|---|---|---|---|---|---|---|---|---|---|---|
| Proportion of patients in the training set | 0.5 | 0.5 | 0.3 | 0.5 | 0.3 | 0.3 | 0.4 | 0.3 | 0.2 | 0.5 | 0.3 |
| Corresponding number of patients | 38 | 38 | 81 | 42 | 152 | 152 | 115 | 106 | 97 | 242 | 160 |

**Supplementary Tab. S3. Minimum Proportion of patients needed in the training dataset without degradation of the C-index and the IBS and corresponding number of patients for the 11 investigated cancers for miRNA-seq data and the Cox model (A), miRNA-seq data and random survival forest (B), mRNA-seq data and the Cox model (C), and mRNA-seq data and random survival forest (D).**

### Supplementary Figures

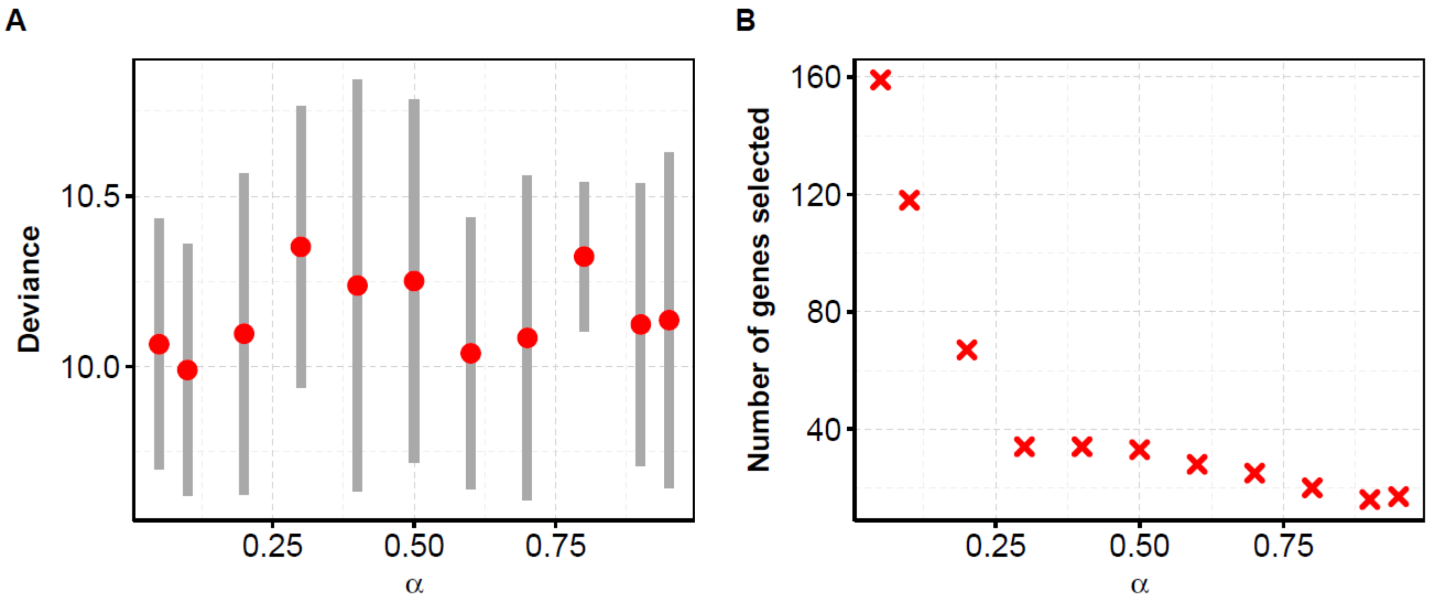


**Supplementary Fig. S1. Deviance and number of genes selected for different values of α for KIRP.**

We computed the deviance by K-fold cross validation (K=5) for each value of α. Similar behavior is observed for the other cancers (data not shown).

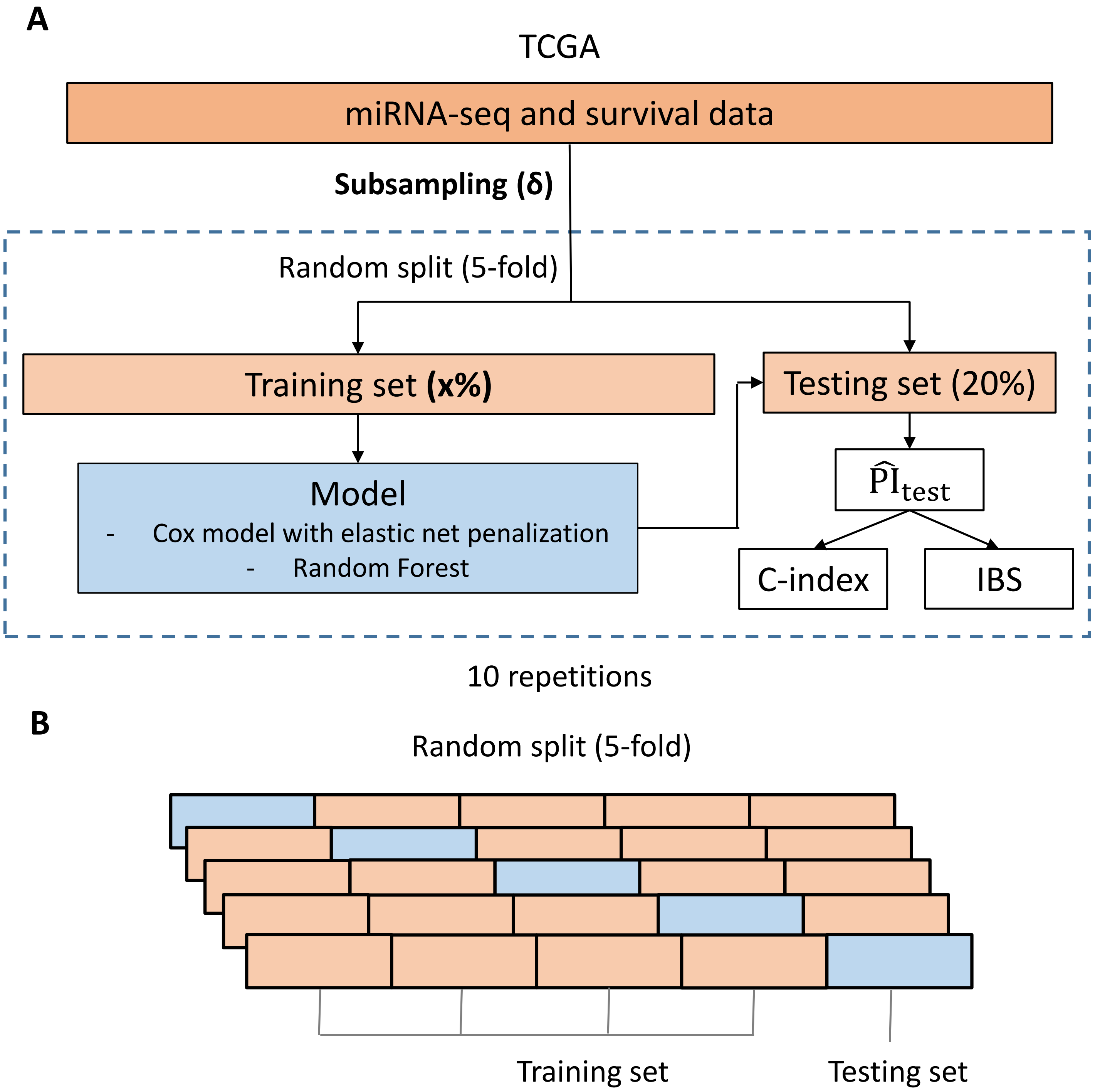


**Supplementary Fig. S2. Procedure for the evaluation of prediction performances.**

CPM corresponds to Count Per Million normalization, RS means `Risk Score' and IBS refers to integrated Brier score.

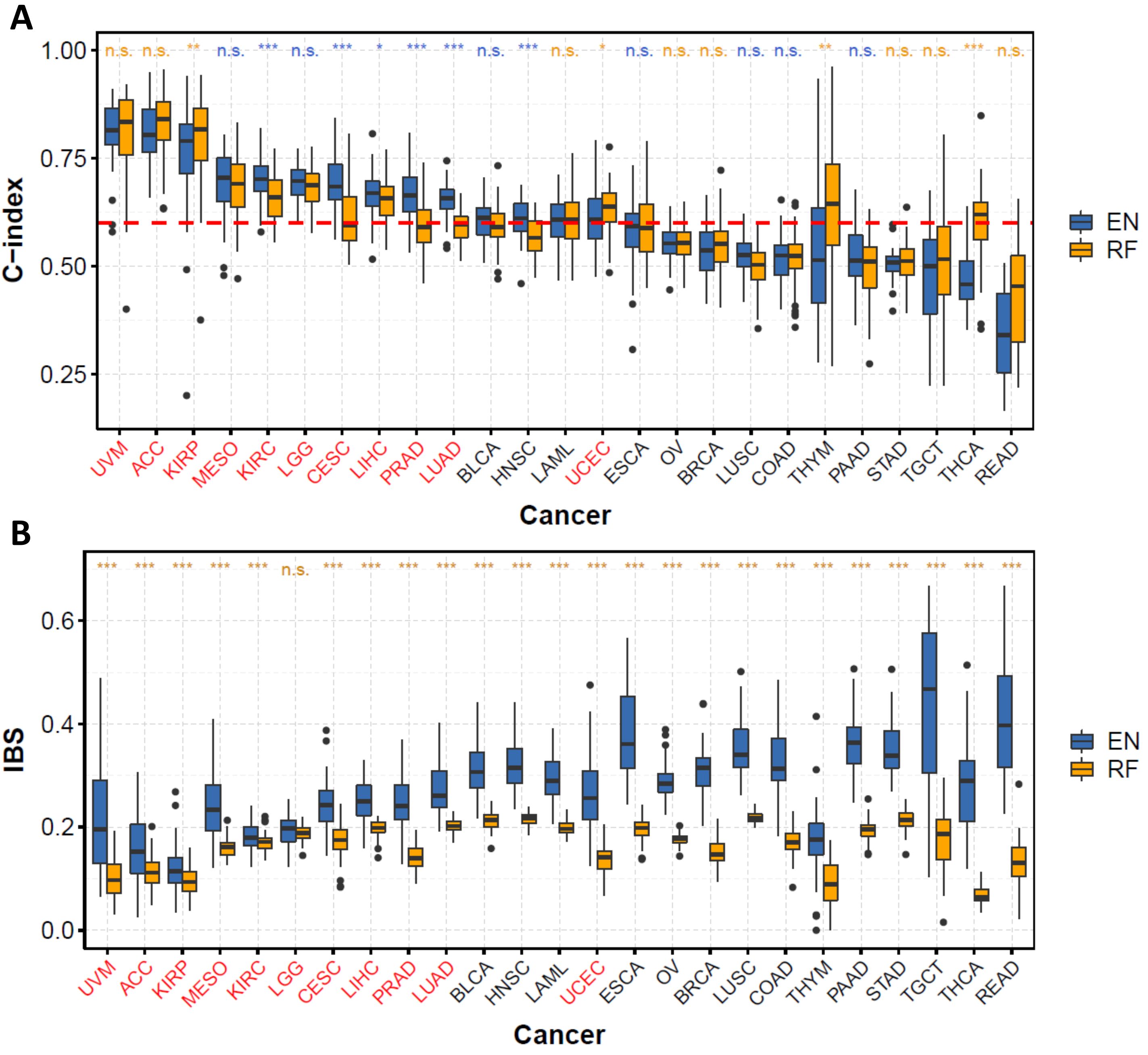


**Supplementary Fig. S3. Boxplot of the C-indices (A) and of the IBS (B) for the Cox model with elastic net penalty (blue) and random forest (orange).**
We computed the metrics by 10 repetitions of a K-fold cross validation (K=5) for all the 25 cancers. We retained 11 cancers (red) that have a median C-index significantly above 0.6 according to a one-sided Wilcoxon test at level 0.05. We corrected the p-values with the Benjamini-Hochberg method.

To compare the predictions obtained with Cox model and random forest, we did a two-sided wilcoxon signed-rank test between C-indices (resp. IBS). Significance level are above each graphics (blue : median C-index is higher or IBS is lower for the Cox model, orange : median C-index is higher or IBS is lower for random forest).

Red dotted horizontal line : C-index of 0.6.
***: p ≤ 0.001, **: p ≤ 0.01, *: p ≤ 0.05, +: p ≤ 0.1, n.s. : p > 0.1

A

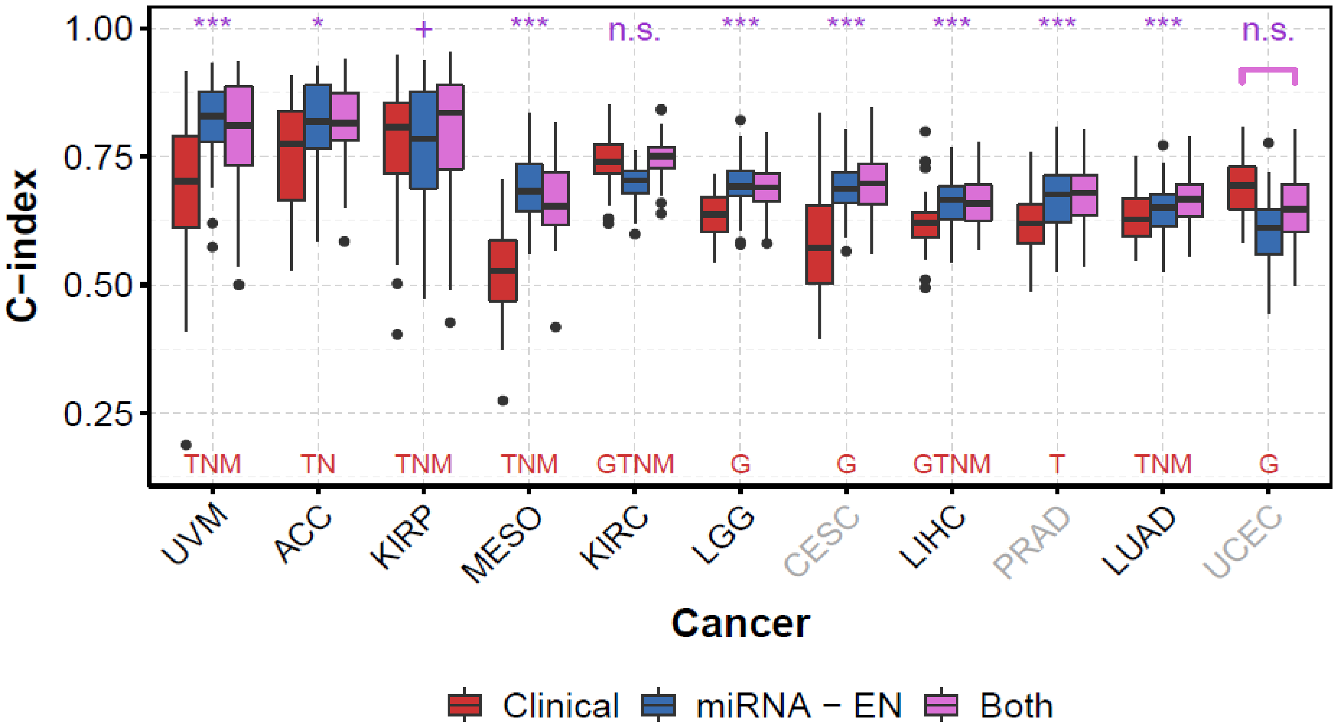


B

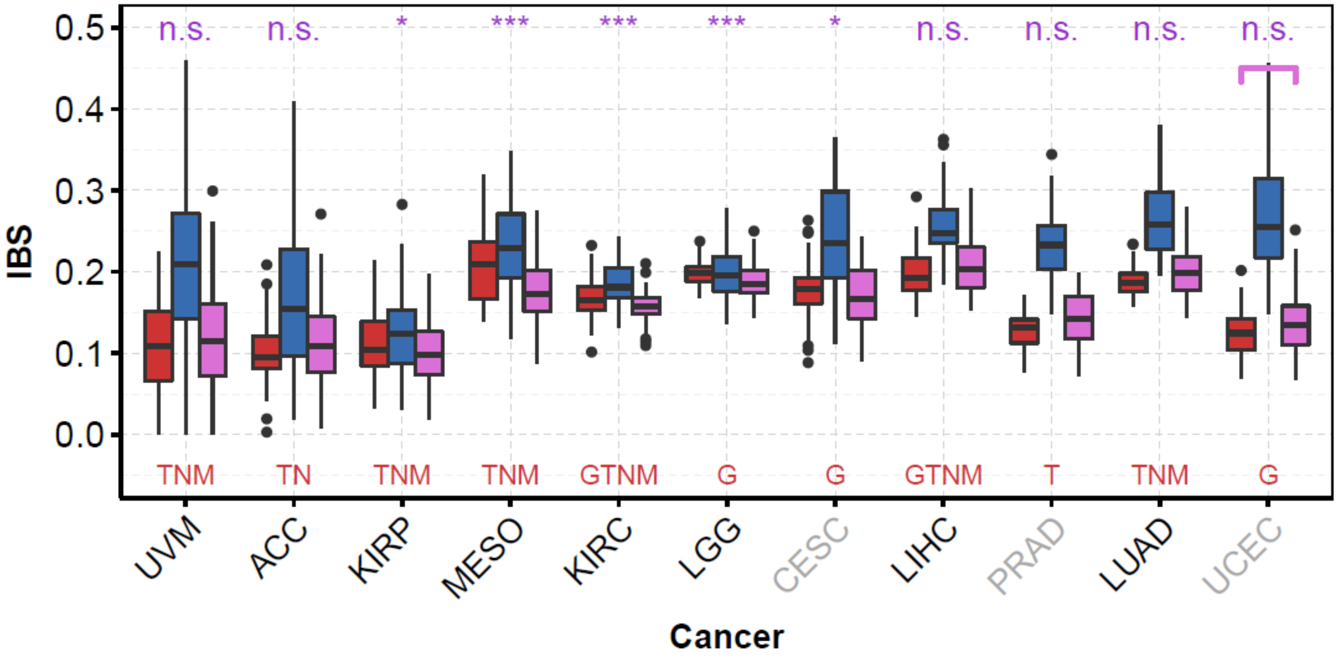


**Supplementary Fig. S4. C-indices (A) and IBS (B) obtained with clinical data alone (red), miRNA-seq data alone (blue), and both clinical and miRNA-seq data (purple) for the 11 cancers investigated and the Cox model with elastic net penalty.**
We computed the metrics by 10 repetitions of a K-fold cross-validation (K=5). We computed p-values of a one-sided Wilcoxon signed-rank test between Clinical and Both Clinical + miRNAs (purple stars at the top of each graphic, Benjamini-Hochberg correction for the 11 p-values). ***: $p \leq 0.001$, **: $p \leq 0.01$, *: $p \leq 0.05$, +: $p \leq 0.1$, n.s. : $p > 0.1$

Red letters at the bottom of each graphics indicate the clinical data available (G: grade; T: tumor; N: node; M: metastasis). Age is available for all cancers, and gender only for non-unisexual cancers (CESC, PRAD, TGCT are sex-specific).

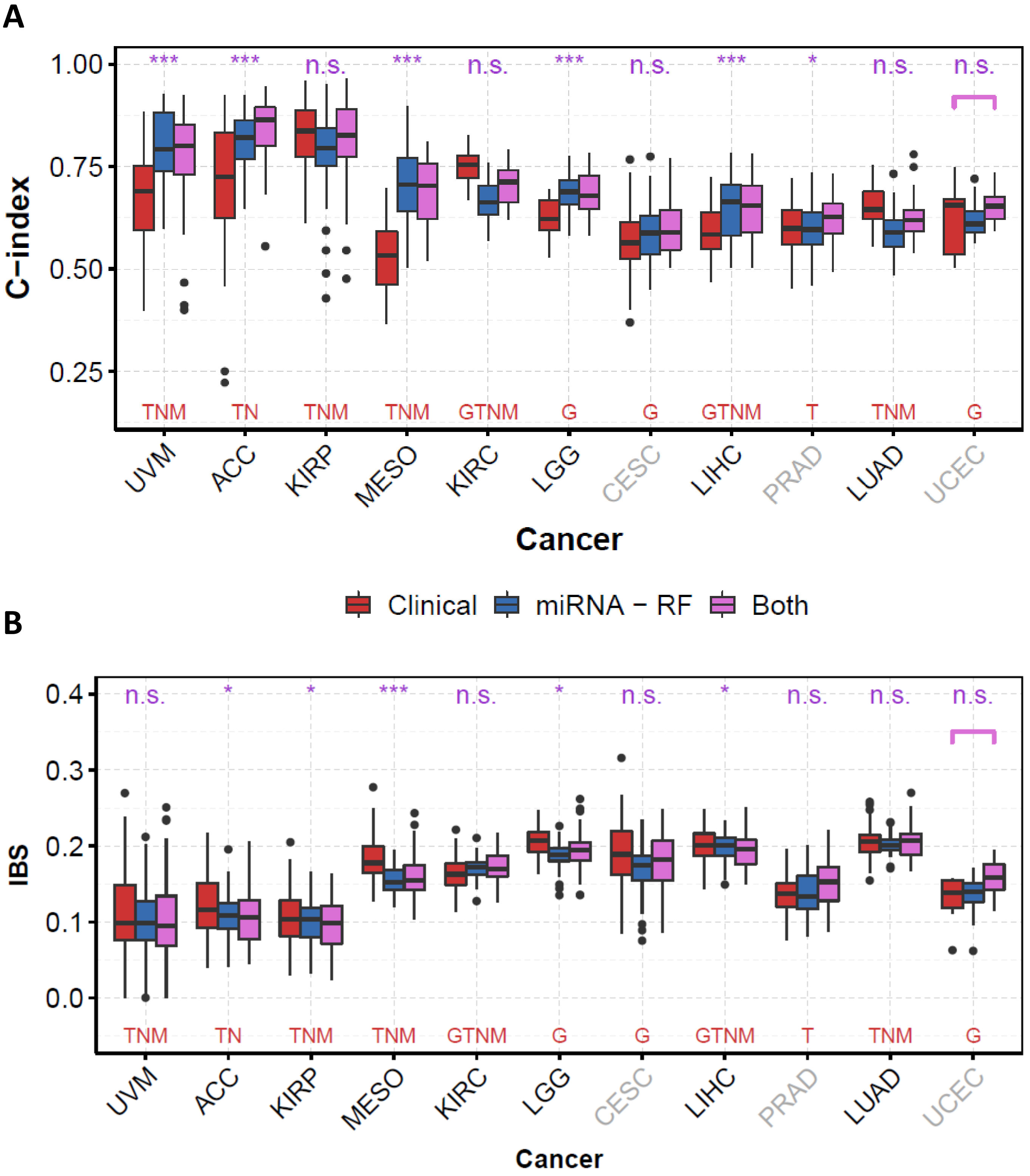


**Supplementary Fig. S5. C-indices (A) and IBS (B) obtained with clinical data alone (red), miRNA-seq data alone (blue), and both clinical and miRNA-seq data (purple) for the 11 cancers investigated and the random survival forest procedure.**

We computed the metrics by 10 repetitions of a K-fold cross-validation (K=5). We computed p-values of a one-sided Wilcoxon signed-rank test between Clinical and Both Clinical + miRNAs (purple stars at the top of each graphic, Benjamini-Hochberg correction for the 11 p-values). ***: $p \leq 0.001$, **: $p \leq 0.01$, *: $p \leq 0.05$, +: $p \leq 0.1$, n.s. : $p > 0.1$

Red letters at the bottom of each graphics indicate the clinical data available (G: grade; T: tumor; N: node; M: metastasis). Age is available for all cancers, and gender only for non-unisexual cancers (CESC, PRAD, TGCT are sex-specific).

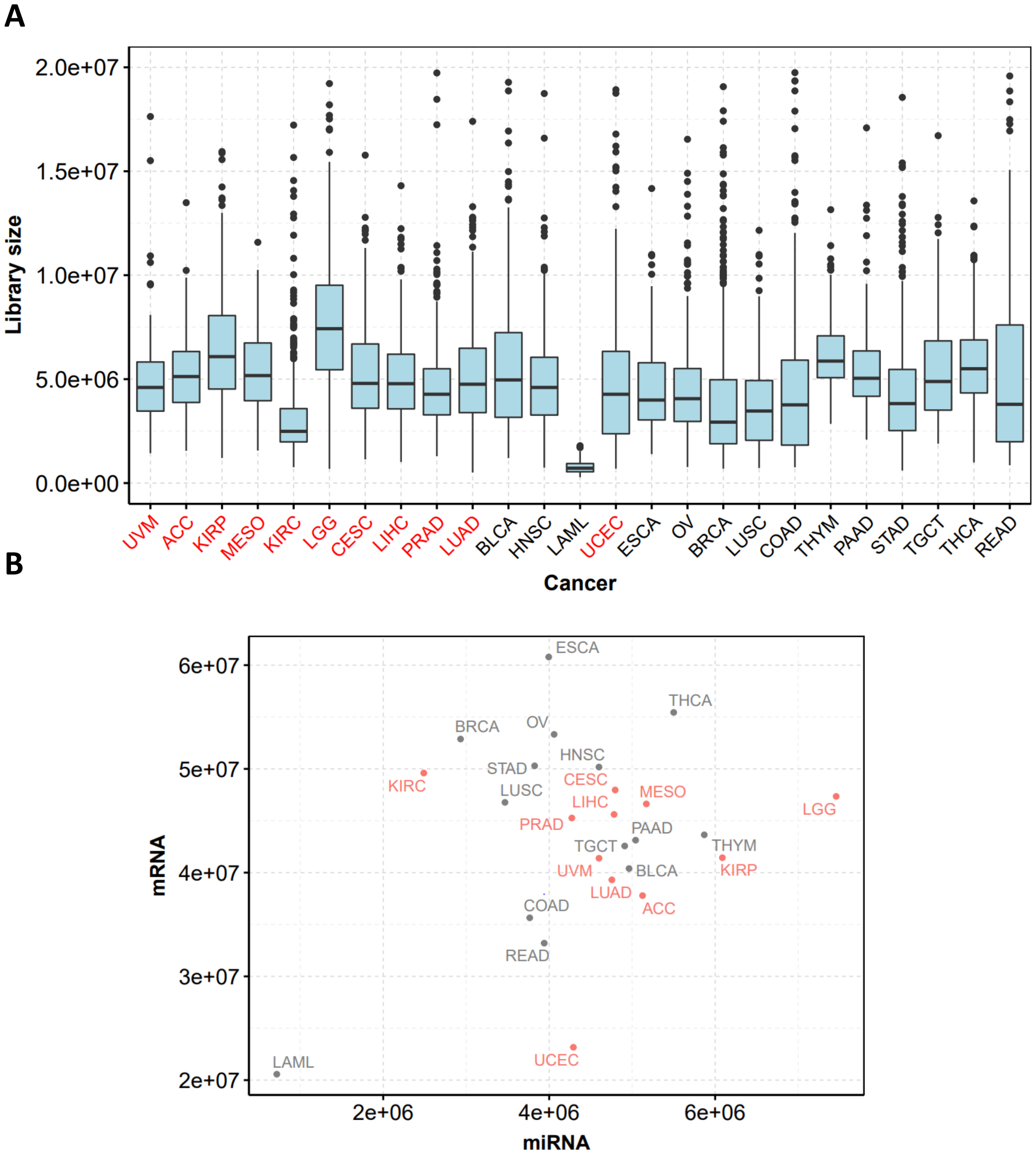


**Supplementary Fig. S6. Distribution of the library size for miRNA-seq data (A), and median library size for mRNA-seq and miRNA-seq data(B) for the 25 cancers of TCGA.**

The 11 cancers investigated for subsampling are in red.

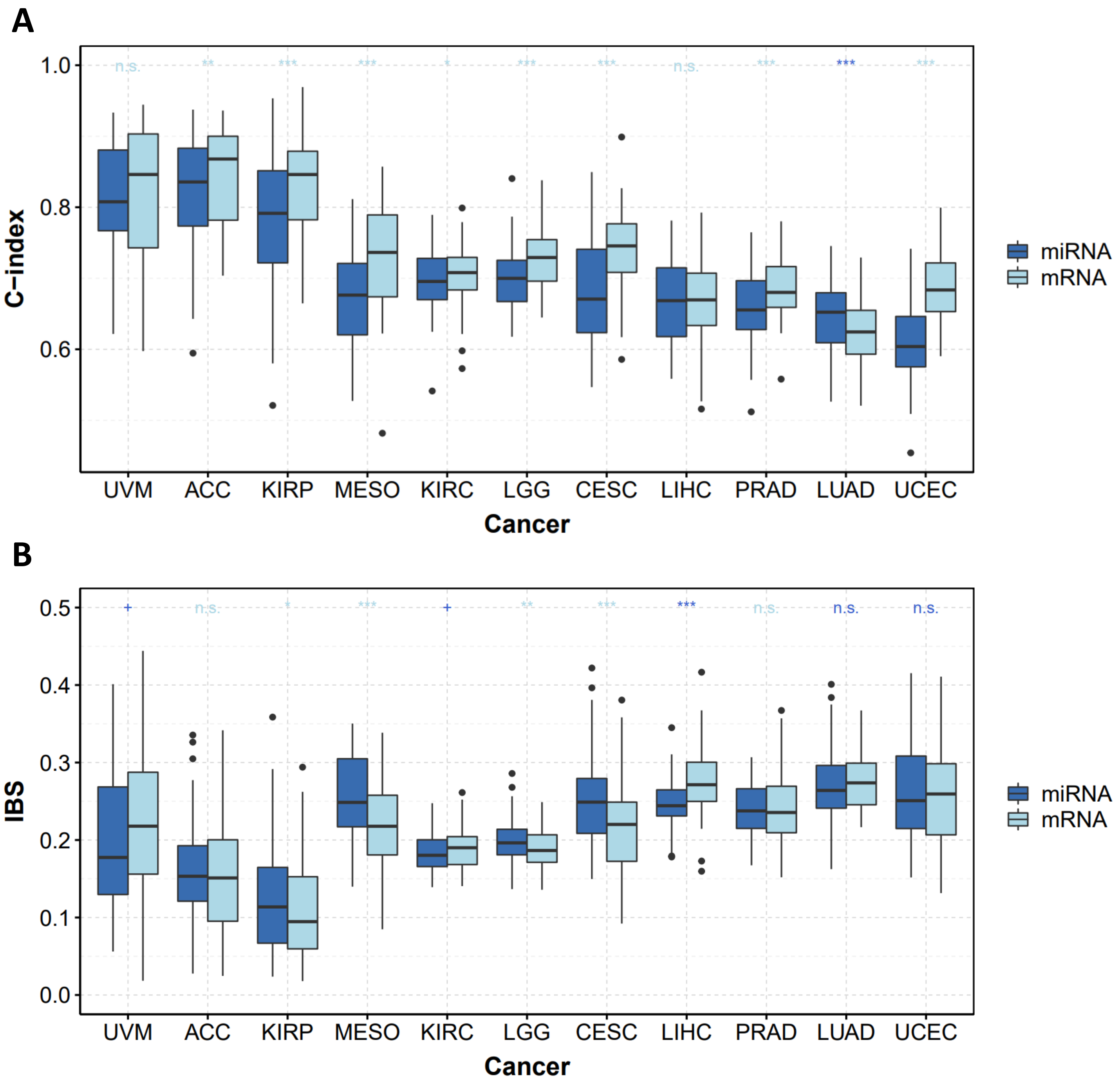


**Supplementary Fig. S7. Boxplot of the C-indices (A) and of the IBS (B) for the Cox model with elastic net penalty for miRNA-seq (blue) and mRNA-seq (lightblue) data.**

We computed  the metrics by 10 repetitions of a K-fold cross validation (K=5) for all the 11 cancers.

To compare the prediction obtained with miRNA-seq and mRNA-seq data, we did a wilcoxon signed-rank test between C-indices (resp. IBS). Significance level with Benjamini-Hocberg correction are above each graphics (blue: median C-index (resp. IBS) is higher (resp. lower) for miRNA-seq data, lightblue: median C-index (resp. IBS) is higher (resp. lower) for mRNA-seq data).

***: $p \leq 0.001$, **: $p \leq 0.01$, *: $p \leq 0.05$, +: $p \leq 0.1$, n.s. : $p > 0.1$

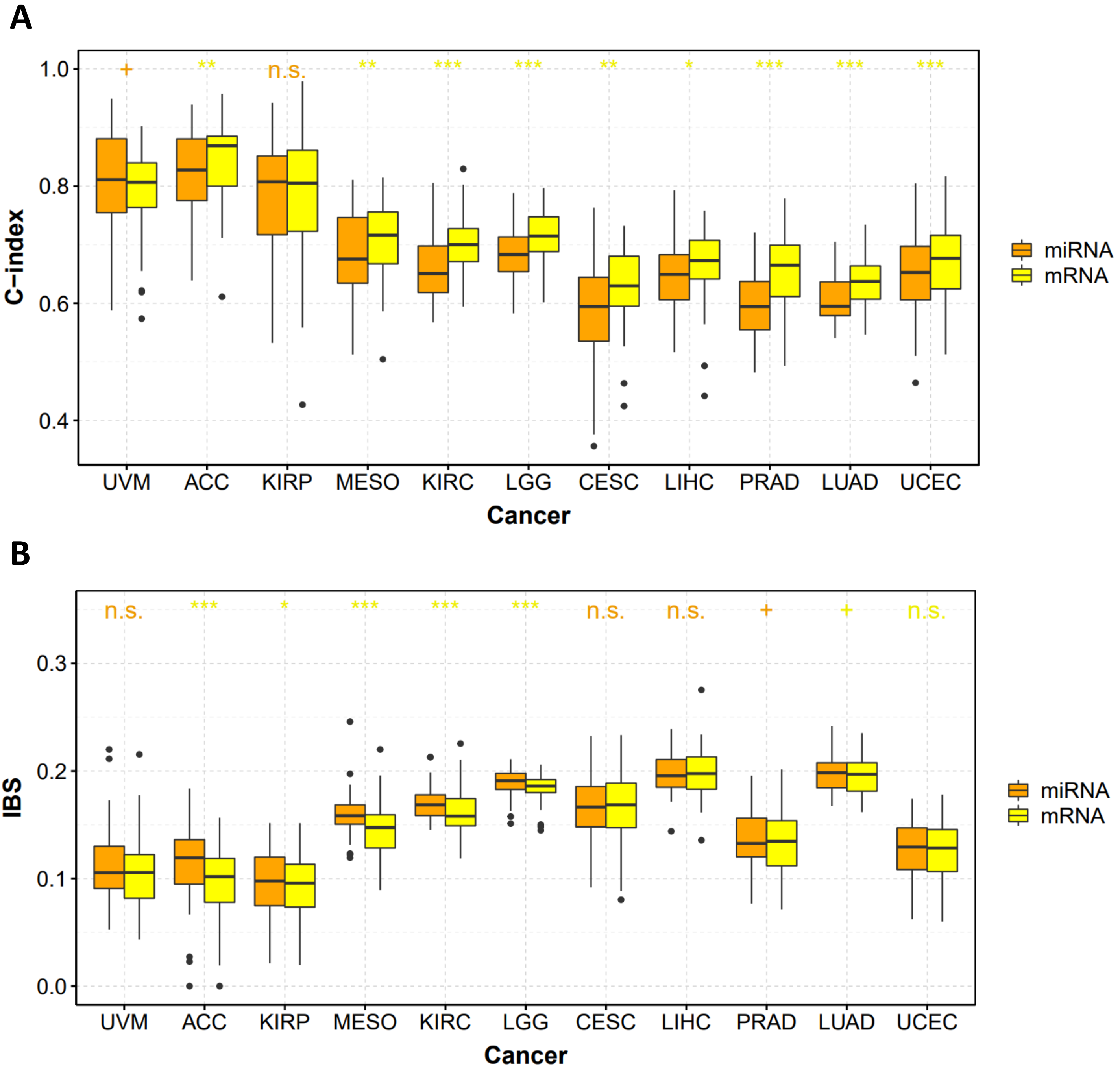


**Supplementary Fig. S8. Boxplot of the C-indices (A) and of the IBS (B) for random survival forest for miRNA-seq (orange) and mRNA-seq (yellow) data.**

We computed the metrics by 10 repetitions of a K-fold cross validation (K=5) for all the 11 cancers.

To compare the prediction obtained with miRNA-seq and mRNA-seq data, we did a wilcoxon signed-rank test between C-indices (resp. IBS). Significance level with Benjamini-Hocberg correction are above each graphics (orange: median C-index (resp. IBS) is higher (resp. lower) for miRNA-seq data, yellow: median C-index (resp. IBS) is higher (resp. lower) for mRNA-seq data).

***: $p \leq 0.001$, **: $p \leq 0.01$, *: $p \leq 0.05$, +: $p \leq 0.1$, n.s. : $p > 0.1$

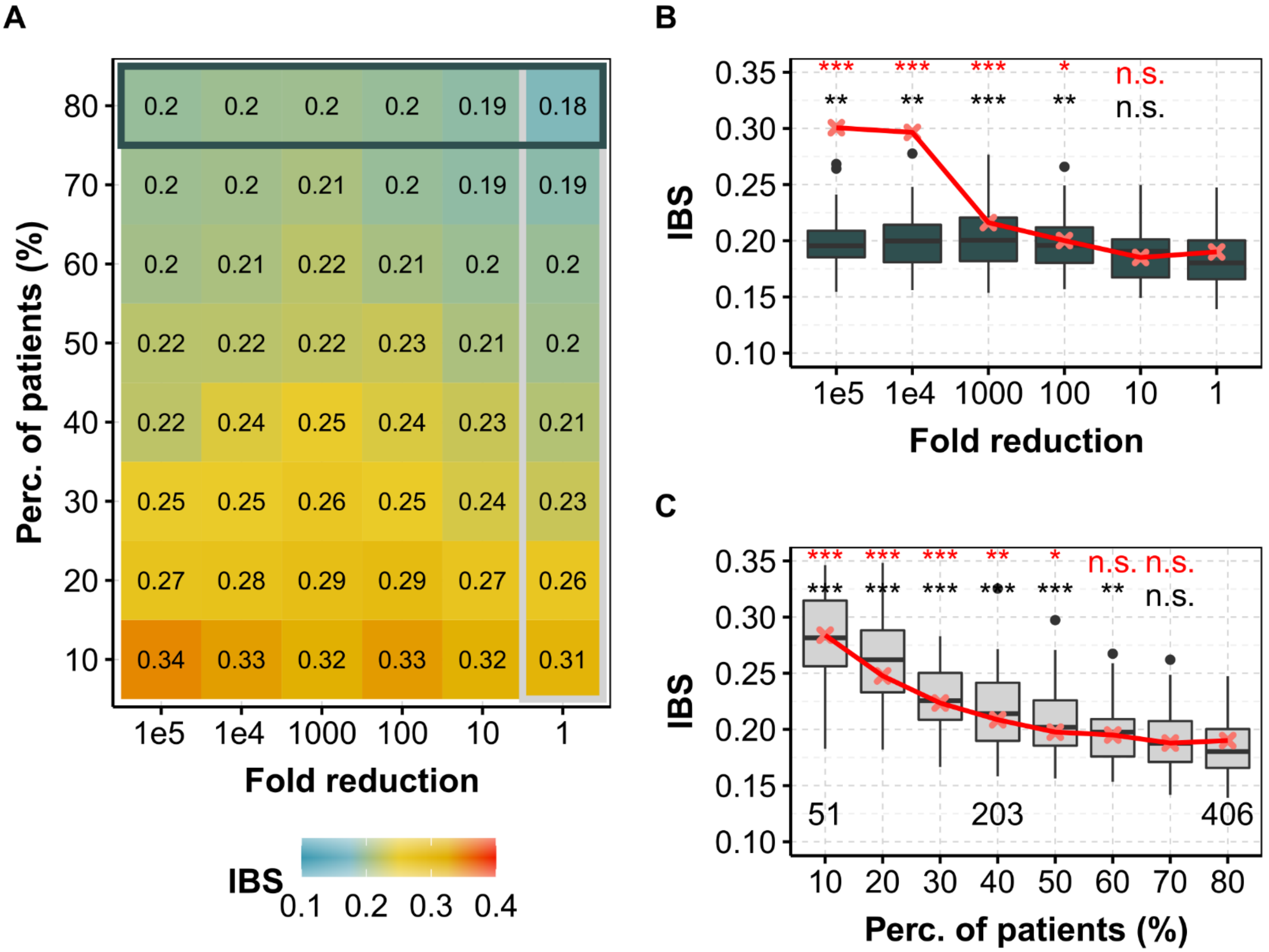


**Supplementary Fig. S9. IBS obtained for different fold reduction factors and percentage of patients in the training dataset for KIRC (ccRCC, TCGA) with the Cox model.**
Same as Fig. 1 but for IBS.

**(A)** Median IBS for different degradation of both sequencing depth (x axis) and percentage of patients (y axis) in the training dataset for miRNA-seq data. Horizontal box highlights the case where all of the 80% of patients are used and corresponds to (B), whereas vertical box focuses on the full available library size and corresponds to (C). **(B)** IBS for different fold reduction factors for miRNA-seq (gray boxplots) and mRNA-seq data (median values, in red) with 80% of the patients in the training dataset. Above is the p-value of a one-sided Wilcoxon test compared to no subsampling (*i.e.* $\delta = 1$). **(C)** IBS for different percentage of patients in the training dataset for miRNA-seq (light gray boxplots) and mRNA-seq data (median values, in red) with original TCGA sequencing depth. Above is the p-value of a one-sided Wilcoxon test compared to full dataset (i.e. 80%).
**red, mRNA-seq; gray (boxplots), miRNA-seq**.

In each case, we computed the IBS by 10 repetitions of a 5-fold cross validation.
***: p ≤ 0.001, **: p ≤ 0.01, *: p ≤ 0.05, +: p ≤ 0.1, n.s. : p > 0.1

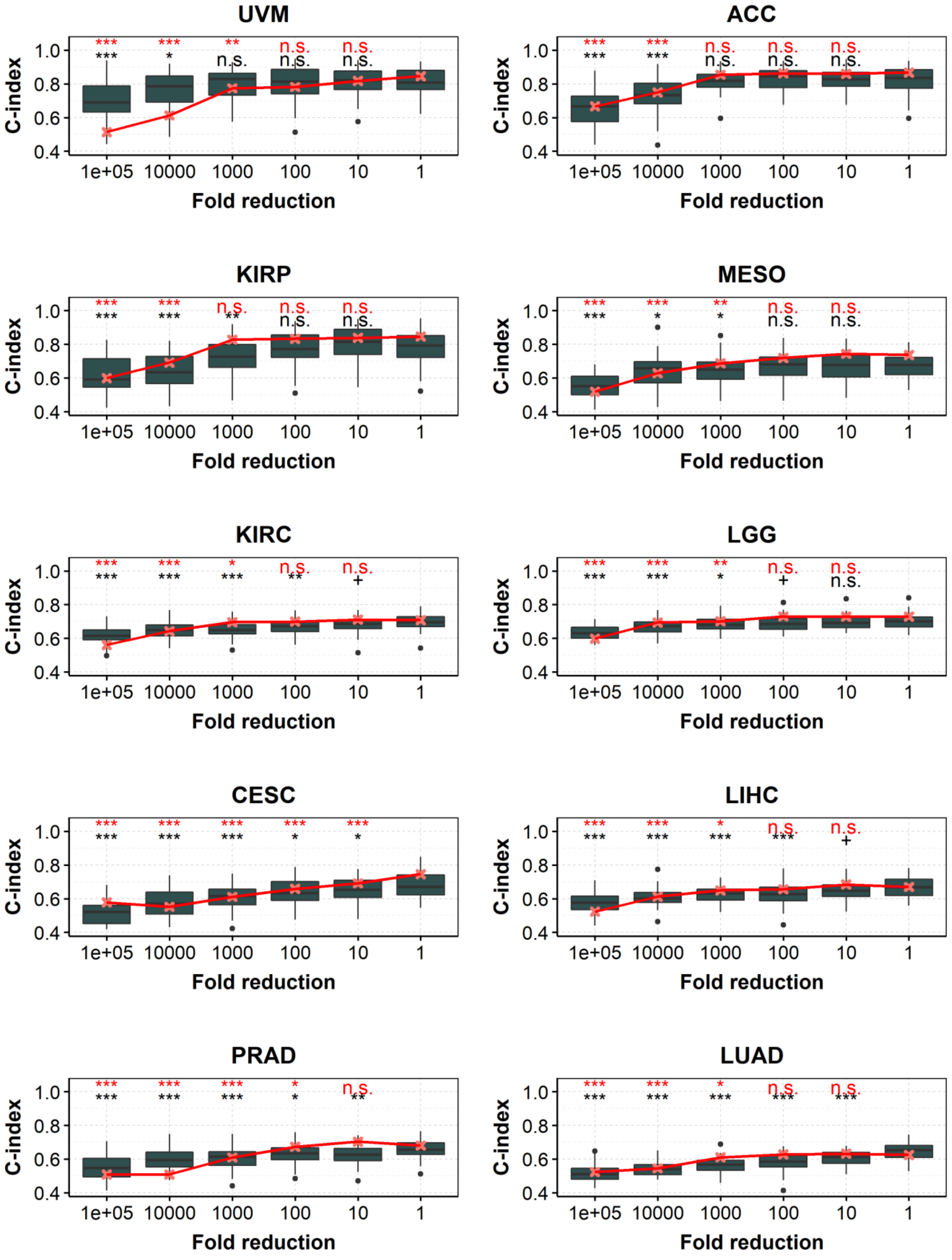


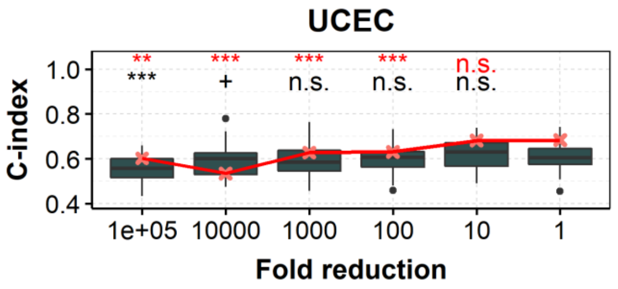


**Supplementary Fig. S10. Corresponds to Figure 1B, but for the 11 investigated cancers. Cox model.**

Full legend next page.

**Supplementary Fig. S10. Distribution of C-indices obtained with the Cox model after fold reduction of miRNA-seq data or mRNA-seq data for the 11 investigated cancers.**
C-index for different fold reduction factors for miRNA-seq (gray boxplots) and mRNA-seq data (median values, in red) with 80% of the patients in the training dataset. Above is the pvalue
of a one-sided Wilcoxon test compared to no subsampling (i.e. d = 1). red, mRNA-seq; gray, miRNA-seq
In each case, we computed the C-indices by 10 repetitions of a 5-fold cross validation.
***: $p \leq 0.001$, **: $p \leq 0.01$, *: $p \leq 0.05$, n.s. : $p > 0.1$.

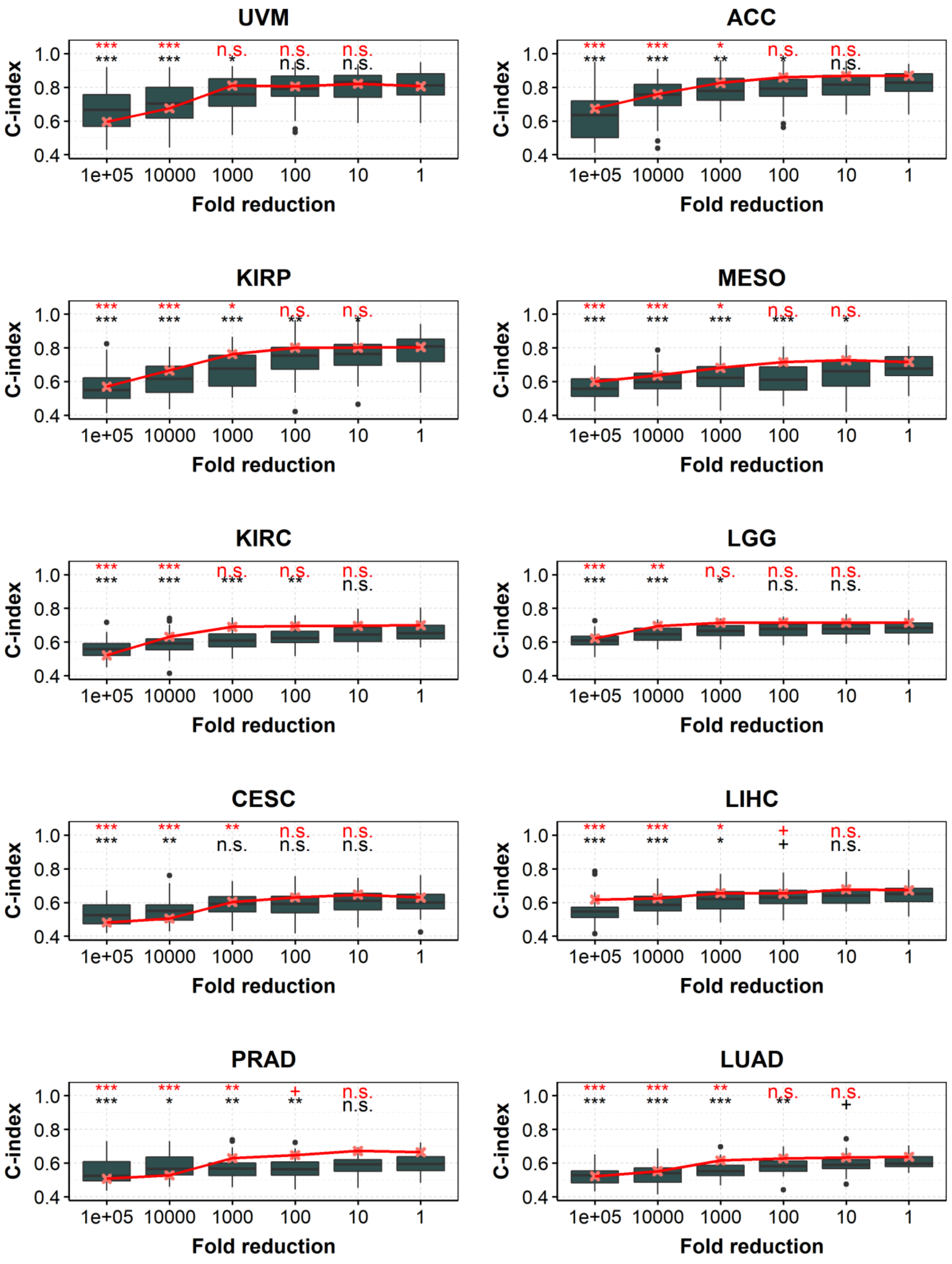


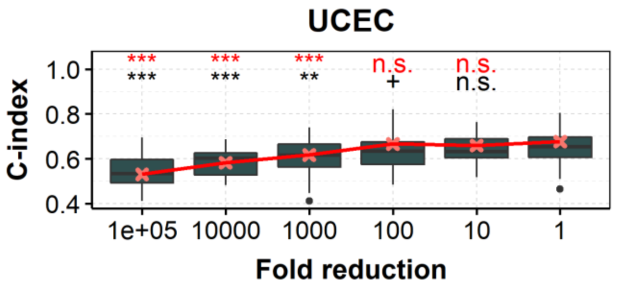


**Supplementary Fig. S11. Corresponds to Figure 1B, but for the 11 investigated cancers and for random survival forest.**
Full legend next page.

**Supplementary Fig. S11. Distribution of C-indices obtained with the random survival forest model after fold reduction of miRNA-seq data or mRNA-seq data for the 11 investigated cancers.**

C-index for different fold reduction factors for miRNA-seq (gray boxplots) and mRNA-seq data (median values, in red) with 80% of the patients in the training dataset. Above is the pvalue
of a one-sided Wilcoxon test compared to no subsampling (i.e. d = 1). red, mRNA-seq; gray, miRNA-seq
In each case, we computed the C-indices by 10 repetitions of a 5-fold cross validation.
***: $p \leq 0.001$, **: $p \leq 0.01$, *: $p \leq 0.05$, n.s. : $p > 0.1$.

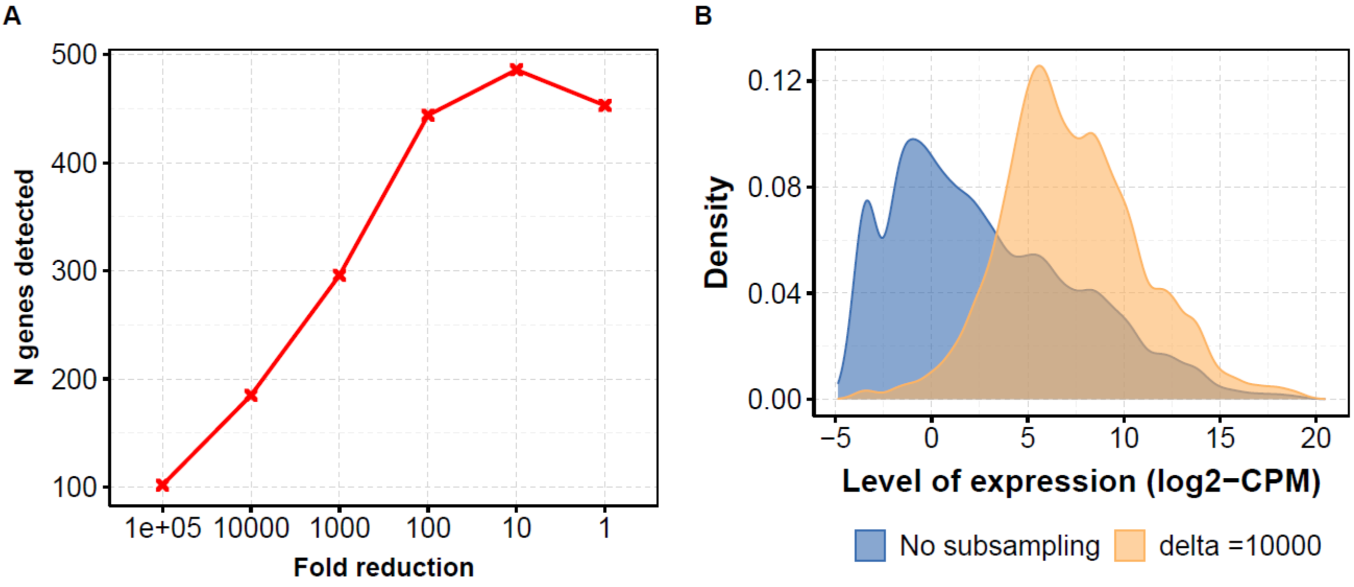


**Supplementary Fig. S12. Number of genes (miRNAs) detected for different fold reduction (A) and level of expression (log2-CPM) of the genes detected without subsampling (blue) and after subsampling by a factor 10,000 for KIRP.**

We defined the 'genes detected' as the miRNAs for which the count per million (CPM) data is higher than 1 for at least 1% of the patients.

Similar results are observed for other cancers and for mRNA-seq data (data not shown).

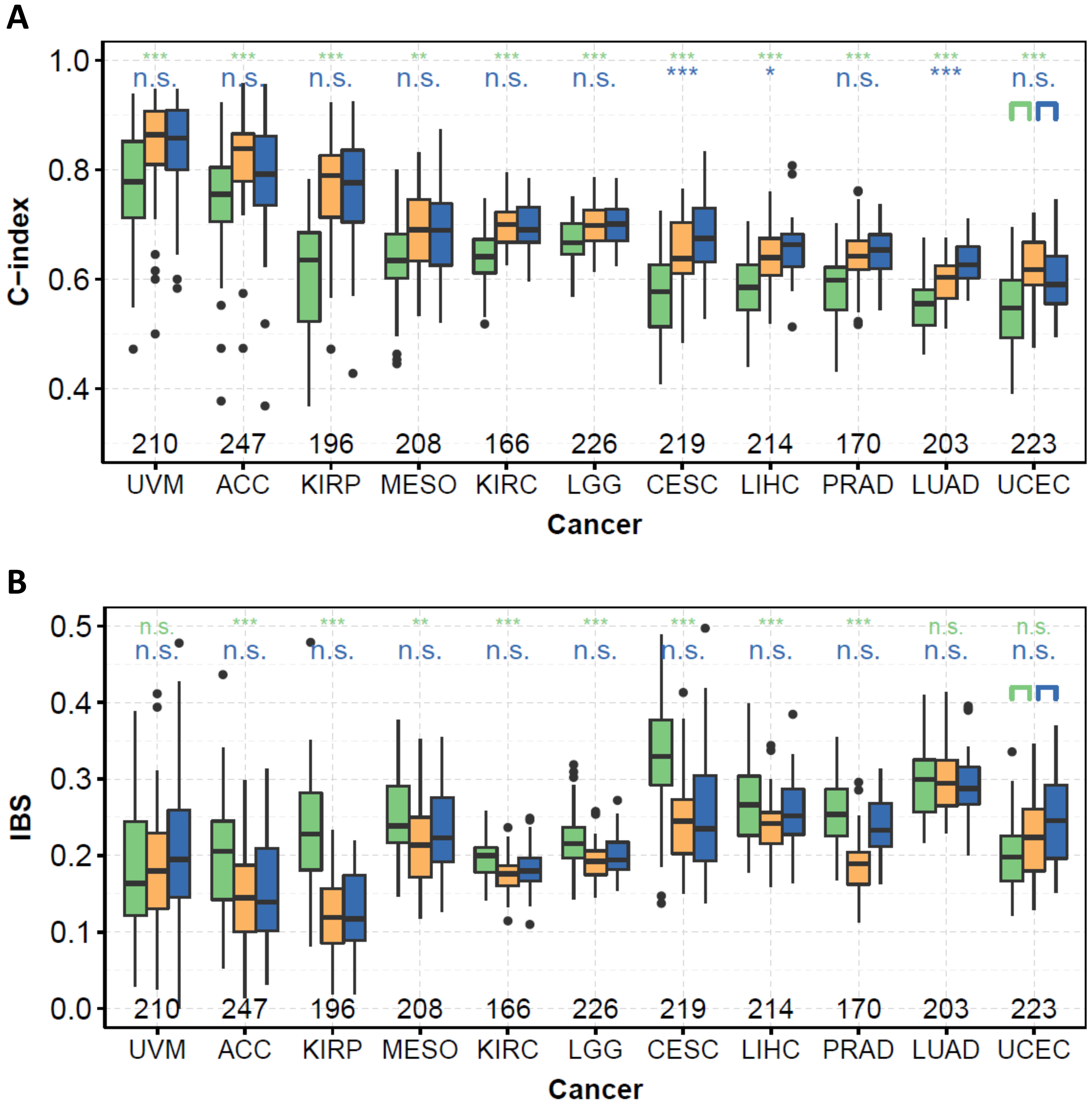


**Supplementary Fig. S13. C-indices (A) and IBS (B) obtained after subsampling by a factor 10,000 (green), with the same miRNAs (~200 most expressed) but without subsampling (orange), and with all the miRNAs (~500) and without subsampling (blue) for the Cox model with elastic net penalty.**

We computed the metrics by 10 repetitions of a K-fold cross-validation (K=5).

Above are the Benjamini-Hochberg corrected indication of the p-values obtained after a pairwise one-sided Wilcoxon test: green stars for orange versus green "scenario", and blue stars for blue versus orange "scenario".

***: $p \leq 0.001$, **: $p \leq 0.01$, *: $p \leq 0.05$, +: $p \leq 0.1$, n.s. : $p > 0.1$

We indicated the number of genes detected after subsampling by a factor 10,000 in black at the bottom of each graphics).

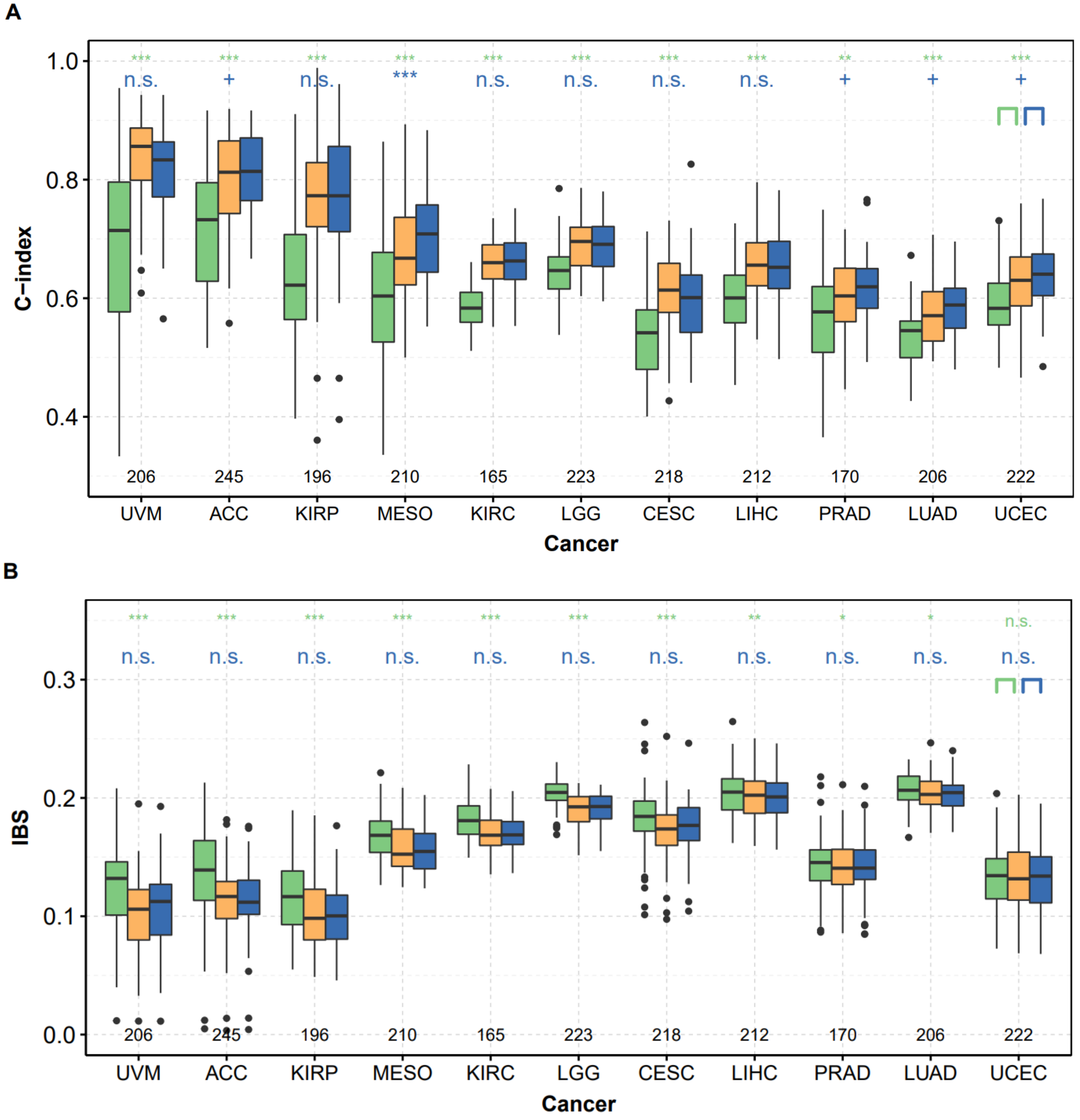


**Supplementary Fig. S14. C-indices (A) and IBS (B) obtained after subsampling by a factor 10,000 (green), with the same miRNAs (~200 most expressed) but without subsampling (orange), and with all the miRNAs (~500) and without subsampling (blue) for random survival forest.**

We computed the metrics by 10 repetitions of a K-fold cross-validation (K=5).

Above are the Benjamini-Hochberg corrected indication of the p-values obtained after a pairwise one-sided Wilcoxon test: green stars for orange versus green "scenario", and blue stars for blue versus orange "scenario".

***: $p \le 0.001$, **: $p \le 0.01$, *: $p \le 0.05$, +: $p \le 0.1$, n.s. : $p > 0.1$

We indicated the number of genes detected after subsampling by a factor 10,000 in black at the bottom of each graphics).

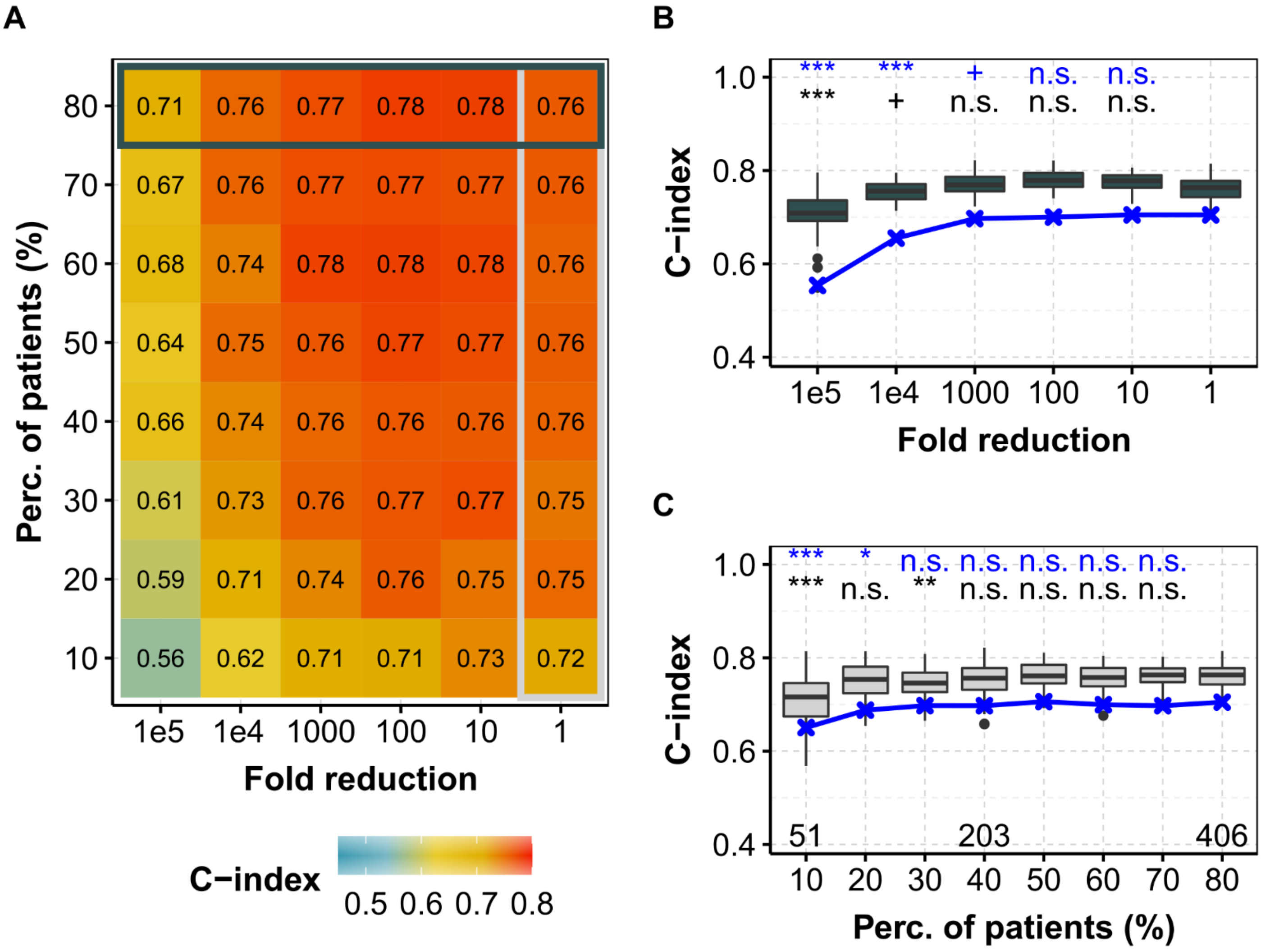


**Supplementary Fig. S15. C-index obtained for different fold reduction factors and percentage of patients in the training dataset for KIRC (ccRCC, TCGA) with the Cox model, assessed on the E-MTAB-1980 dataset.**
Same as Fig. 1 but for the independent E-MTAB-1980 dataset.

**(A)** Median C-index for different degradation of both sequencing depth (x axis) and percentage of patients (y axis) in the training dataset for miRNA-seq data. Horizontal box highlights the case where all of the 80% of patients are used and corresponds to (B), whereas vertical box focuses on the full available library size and corresponds to (C). **(B)** C-index for different fold reduction factors for miRNA-seq (gray boxplots) and mRNA-seq data (median values, in red) with 80% of the patients in the training dataset. Above is the p-value of a one-sided Wilcoxon test compared to no subsampling (*i.e.* δ = 1). **(C)** C-index for different percentage of patients in the training dataset for miRNA-seq (light gray boxplots) and mRNA-seq data (median values, in red) with original TCGA sequencing depth. Above is the p-value of a one-sided Wilcoxon test compared to full dataset (i.e. 80%).
**blue, mRNA-seq from TCGA; gray (boxplots), mRNA-seq from E-MTAB-1980**.

In each case, we computed the C-indices by 10 repetitions of a 5-fold cross validation.
***: $p \leq 0.001$, **: $p \leq 0.01$, *: $p \leq 0.05$, +: $p \leq 0.1$, n.s. : $p > 0.1$

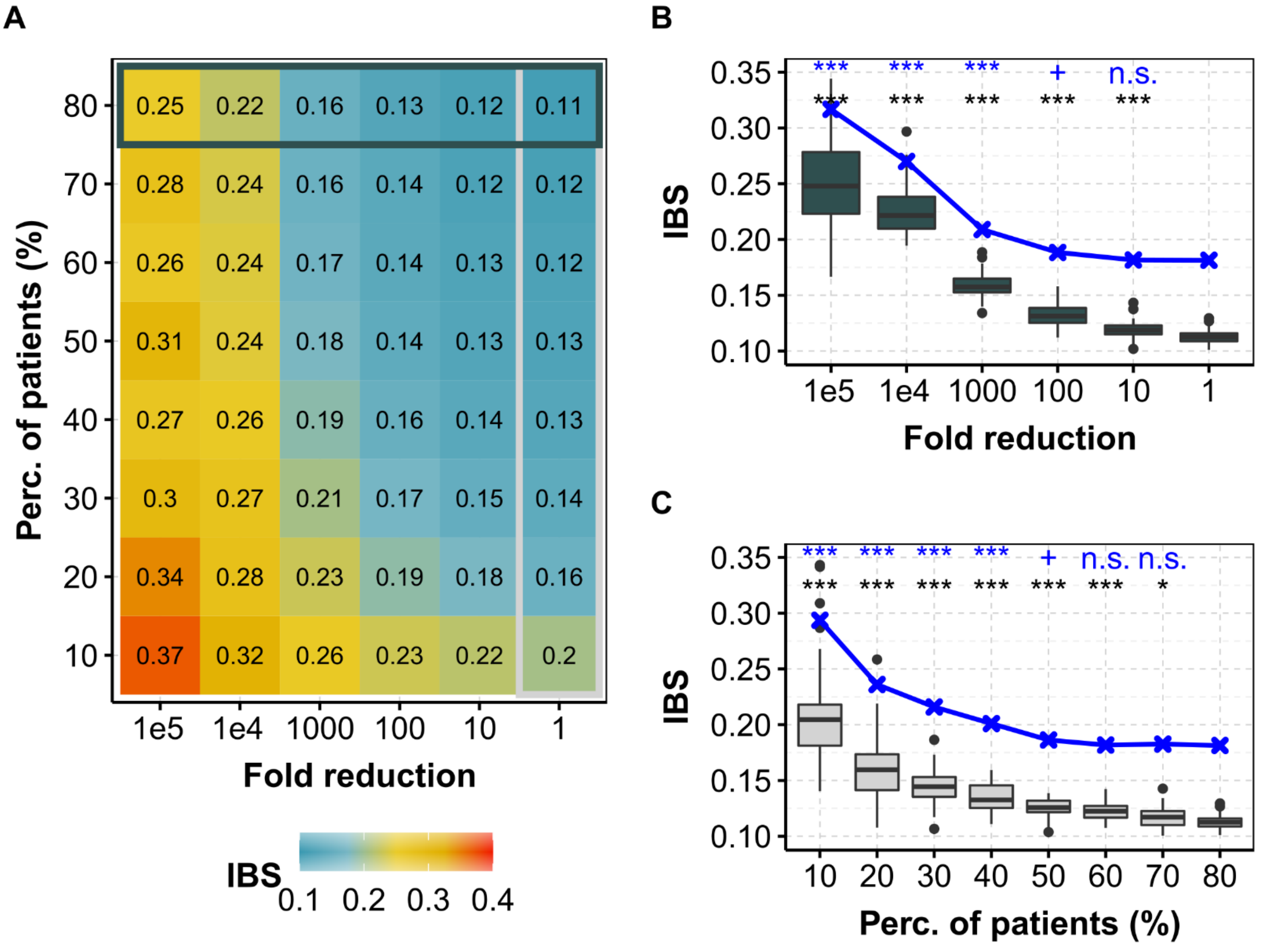


**Supplementary Fig. S15. IBS obtained for different fold reduction factors and percentage of patients in the training dataset for KIRC (ccRCC, TCGA) with the Cox model, assessed on the E-MTAB-1980 dataset.**
Same as Fig. 1 but for the independent E-MTAB-1980 dataset and for IBS.

**(A)** Median IBS for different degradation of both sequencing depth (x axis) and percentage of patients (y axis) in the training dataset for miRNA-seq data. Horizontal box highlights the case where all of the 80% of patients are used and corresponds to (B), whereas vertical box focuses on the full available library size and corresponds to (C). **(B)** IBS for different fold reduction factors for miRNA-seq (gray boxplots) and mRNA-seq data (median values, in red) with 80% of the patients in the training dataset. Above is the p-value of a one-sided Wilcoxon test compared to no subsampling (*i.e.* $\delta = 1$). **(C)** IBS for different percentage of patients in the training dataset for miRNA-seq (light gray boxplots) and mRNA-seq data (median values, in red) with original TCGA sequencing depth. Above is the p-value of a one-sided Wilcoxon test compared to full dataset (i.e. 80%).
**blue, mRNA-seq from TCGA; gray (boxplots), mRNA-seq from E-MTAB-1980**.

In each case, we computed the IBS values by 10 repetitions of a 5-fold cross validation.
***: $p \leq 0.001$, **: $p \leq 0.01$, *: $p \leq 0.05$, +: $p \leq 0.1$, n.s. : $p > 0.1$
